# Spermatogonia derived RpS13 non-autonomously regulates the cyst cell differentiation via Rho1-mediated adhesion junctions in *Drosophila* testis

**DOI:** 10.1101/2025.03.25.645343

**Authors:** Min Wang, Li Zhang, Jianbo Xu, Yanxin Zhang, Min Jiang, Jingzhe Song, Xilong Liu, Lingling Gao, Dan Lu

## Abstract

**Objective:** In the testis of *Drosophila*, spermatogonia transit-amplification (TA) divisions are necessary for germline proliferation and differentiation. Both the germline and the surrounding somatic cyst cells (SCCs) have a significant influence on these processes. However, the underlying mechanism is largely unknown.

**Methods:** Flies were used to generate tissue-specific gene knockdown. Immunofluorescence was used to determine protein localization and expression pattern. Small interfering RNAs were used to knockdown genes in S2 cells. qRT-PCR was used to examine the relative mRNA expression level.

**Results:** Our findings indicate that spermatogonial RpS13 regulates cyst cell growth through a non-autonomous manner. In terms of mechanism, RpS13 may regulate the adhesion junctions of the soma–germline via the DE-cad and Arm proteins in addition to dpERK, which is found in SCCs. Interestingly, Rho1 and RpS13 mutually inhibit each other in *Drosophila* S2 cells. Additionally, Rho1 controlled cell adhesions that rely on DE-cad, Arm, and dpERK and imitated the actions of RpS13 in *Drosophila* testis.

**Conclusion:** All of these findings implied that during spermatogonia TA-divisions in the testis of *Drosophila*, RpS13 regulates the development of cyst cells via Rho1-mediated adhesion junctions.

## Introduction

In the context of tissue regeneration and homeostasis, the precise generation and differentiation of stem cell progeny are essential. In *Drosophila* melanogaster testes, this process involves the coordinated asymmetric division of a germline stem cell (GSC) and two neighboring somatic cyst stem cells (CySCs), yielding a gonialblast and two SCCs, respectively [1,2]. A spermatogonial cyst is subsequently formed when a gonialblast is enveloped by two SCCs [2]. This intricate process is regulated by multiple integrated signals that maintain homeostasis and ensure proper soma–germline interactions [3]. Notably, EGFR inactivation in the adjacent SCCs inhibits the differentiation of germline cells during the spermatogonia transit-amplifying (TA) division phase in *Drosophila* testis [4]. Additionally, the synchronization of germline divisions in the *Drosophila* testis relies on somatic ERK activation during the TA phase [5]. Furthermore, somatic Rac1 and Cdc42, which are downstream effectors of the non-canonical Wnt signaling pathway, play critical roles in regulating germline differentiation and proliferation [6]. EGFR activation, mediated by Rac1, is proposed to facilitate the proper encapsulation of germ cells, a key regulatory step in TA progression [7]. This detailed molecular coordination underscores the complexity of germline–somatic interactions in maintaining testicular homeostasis and ensuring accurate tissue regeneration.

Defects in cell adhesion and signal transduction mechanisms can result in cellular mislocalization, uncontrolled proliferation, and aberrant differentiation, ultimately leading to pathological conditions such as tissue overgrowth, tumorigenesis, and cancer metastasis [8]. Notably, the maintenance of stem cells is critically dependent on adhesion between stem cells and their niche. In *Drosophila* melanogaster, clusters of adhesion junctions have been identified between male GSCs and adjacent hub cells [8]. DE-cadherin-mediated adhesion plays a pivotal role in the initial recruitment and subsequent anchoring of germline-derived stem cells to their niche, as well as in orchestrating cell migration during *Drosophila* oogenesis [9]. Furthermore, the aminopeptidase Sda, which is essential for the accumulation of mature DE-cadherin, is implicated in the maintenance of germline stem cells and spermatogonial dedifferentiation within the testicular niche of *Drosophila* [10]. Immunofluorescence studies have demonstrated that the *Drosophila* E-cadherin homolog Shotgun (Shg) and the β-catenin homolog Armadillo (Arm) are densely localized at the interface between GSCs and hub cells in males [11]. Arm, as a multifunctional protein, not only interacts with cadherin to facilitate cell adhesion but also links to the actin cytoskeleton [12,13]. It is hypothesized that adhesion-mediated cell–cell interactions are crucial for retaining stem cells within their niche, ensuring their proximity to maintenance signals while shielding them from differentiation cues [14]. This intricate interplay of adhesion molecules and signaling pathways underscores the importance of proper cell adhesion in stem cell biology and tissue homeostasis.

Ribosome biogenesis is precisely regulated in a stage-specific manner throughout gametogenesis, reflecting its critical role in reproductive development. A key requirement during oogenesis is the production and accumulation of a substantial reservoir of ribosomes, which are essential to sustain early embryonic development prior to the activation of zygotic transcription [15]. Emerging evidence highlights that ribosome biogenesis is not merely a housekeeping process but serves as a crucial regulatory mechanism in stem cell biology, potentially mediating the transition between stem cell maintenance and differentiation [16]. In *Drosophila* melanogaster, this regulation is exemplified by the specialized function of eRpL22 paralog-specific ribosomes, which modulate the translation of specific mRNAs during spermatogenesis, thereby contributing to the precise control of germ cell development [17]. The ribosome-specific MrpL55 protein is localized in the mitochondria of *Drosophila* S2 cells [18]. In the context of *Drosophila* testis, RpL6 plays a crucial role in regulating the self-renewal and differentiation of GSCs. Notably, S2 cells overexpressing RpL6 exhibit enhanced proliferative capacity and significantly accelerate the process of cell death [19]. Additionally, germ cell-specific expression of RpS5b is essential for proper egg chamber development, as it ensures the homeostasis of functional ribosomes [20]. In the *Drosophila* ovary, ribosomal assembly factors are indispensable for the regulation of stem cell cytokinesis and are critical for facilitating the transition from self-renewal to differentiation [21,22]. These findings underscore the intricate and multifaceted role of ribosome biogenesis in gametogenesis and stem cell regulation, emphasizing its importance in developmental biology.

Our prior research confirmed that RpS13 was required for the self-renewal and differentiation of GSCs in *Drosophila* testes [23]. In our current study, we observed that spermatogonial RpS13 and Rho1 play a non-autonomous role in regulating cyst cell differentiation. Furthermore, these genes influence the expression of adhesion proteins such as DE-cad and Arm, as well as dpERK signaling pathways in the *Drosophila* testis.

## Materials and methods

### Fly strains

All *Drosophila* strains were maintained under controlled conditions at 25 °C with a relative humidity of 40–60%, utilizing a standard cornmeal molasses agar medium as the growth substrate. The transgenic RNA interference (RNAi) lines employed in this study were procured from the TsingHua Fly Center (THFC, Beijing, China). Specifically, the lines used were UAS–RpS13 RNAi (#THU0667) and UAS–Rho1 RNAi (#THU3565). Additionally, the Bam-Gal4; Δ86/+ line was generously provided by Dr. DH Chen from the Institute of Zoology, Chinese Academy of Sciences, Beijing, China.

### Fly cross strategy

Transgenic UAS-RNAi virgin females were randomly chosen and crossed with male flies of the Bam-Gal4; Δ86/+ line. The resulting progeny were reared at a constant temperature of 25°C. Subsequently, F1 males exhibiting the specific genotype (Bam > RNAi or Bam > RNAi; Δ86/+) were identified and selected within a 48-hour window post-eclosion for subsequent experimental procedures. The W1118 and Δ86/+ lines served as the control groups in this study.

### Immunostaining and antibodies

Fly testes were dissected and prepared in 1× phosphate-buffered saline (PBS), followed by fixation in 4% paraformaldehyde for 30 minutes at room temperature. After fixation, the samples were washed three times with 0.3% PBS containing Triton X-100 (PBST) to permeabilize the tissues. Subsequently, the testes were blocked in 5% bovine serum albumin (BSA) for 30 minutes to reduce non-specific binding. The tissues were then incubated overnight at 4°C with primary antibodies diluted in 5% BSA. Following primary antibody incubation, the testes were washed three times with 0.3% PBST to remove unbound antibodies. Secondary antibodies, conjugated with Alexa Fluor 488 (A488) or Cyanine 3 (Cy3) (Molecular Probes and Jackson Immunologicals), were applied at a dilution of 1:1000 and incubated at room temperature for 1 hour in the dark. After three additional washes with 0.3% PBST, the testes were stained with Hoechst-33342 (1.0 mg/ml, C0031, Solarbio, Beijing, China) for 5 minutes to label nuclei. Finally, the samples were mounted in glycerol solution for imaging.

The primary antibodies used in this study included: rat anti-Zfh1 (1:2000, kindly provided by Tong Lab), mouse anti-Eya (1:50, Developmental Studies Hybridoma Bank [DSHB], USA), rat anti-DE-cadherin (1:20, DSHB, USA), mouse anti-Armadillo (1:50, DSHB, USA), rabbit anti-Vasa (1:200, Santa Cruz Biotechnology, USA), mouse anti-Rho1 (1:50, DSHB, USA), and rabbit anti-phospho-ERK (dpERK) (1:200, Cell Signaling Technology [CST], USA).

### Images acquisition and analysis

Confocal images were obtained with the Zeiss LSM800 system (Carl Zeiss, Oberkochen, Germany) and were processed with Adobe Photoshop Software (Adobe, San Jose, CA, USA). Quantification analysis of fluorescence intensity was performed with ImageJ software.

### Cell culture and transfection

*Drosophila* Schneider 2 (S2) cells, procured from the *Drosophila* Genomics Resource Center, were maintained in Schneider’s *Drosophila* Medium (21720024, Gibco, USA) supplemented with 10% heat-inactivated fetal bovine serum (04-001-1ACS; Biological Industries, Israel) and cultured at a constant temperature of 28°C. The cells were passaged every 3-4 days at a 1:4 ratio to ensure optimal growth and viability. For experimental purposes, S2 cells were seeded in 6-well plates and allowed to reach 70%–80% confluence. To perform the knockdown assay, the cells were transfected using Lipofectamine 2000 (Lipo2000; 11668019, Invitrogen, USA). The specific small interfering RNAs (siRNAs) used for knockdown were designed and synthesized by GenePharma (Shanghai, China). Additional details regarding the siRNA sequences and target genes are provided in Supplementary Table S1.

### Quantitative real-time reverse transcription PCR

Total RNA was extracted using TRIzol reagent (9108, Takara, Japan). Subsequently, reverse transcription was carried out following the manufacturer’s protocol with the PrimeScript RT Reagent Kit (RR037A, Takara, Japan). Quantification of gene expression was performed via quantitative PCR (qPCR) using TB Green Premix Ex Taq II (RR420A, Takara, Japan). The qRT-PCR reactions were conducted on an Agilent Mx3000P Real-Time PCR System (Agilent Technologies, Santa Clara, CA, USA). Relative mRNA expression levels were calculated using the 2-^△ △ Ct^ method, where Ct values represent threshold cycles. For normalization, glyceraldehyde-3-phosphate dehydrogenase (GAPDH) was employed as the internal reference gene. The primer sequences used in this study are detailed in Supplementary Table S2.

### Statistical analysis

All experiments were independently repeated a minimum of three times to ensure reproducibility. Quantitative data are presented as the mean ± standard error of the mean (SEM). Statistical analyses, including Student’s t-test and one-way ANOVA, were performed using GraphPad Software (La Jolla, CA, USA). A *P*-value of less than 0.05 (*P* < 0.05) was considered statistically significant. This rigorous approach ensured the reliability and validity of the experimental results.

## Results

### Spermatogonia derived RpS13 non-autonomously regulate the cyst cell differentiation

The function of RpS13 in *Drosophila* testes was investigated through an RNA interference (RNAi) assay driven by the Bam-Gal4 system. Germ cell differentiation in *Drosophila* testes is regulated by the Bam protein, a key differentiation factor, as demonstrated in previous studies [24, 25]. To enhance the phenotypic characterization of germ cell differentiation, the heterozygous bam^Δ86^ null allele (Δ86/+) [26] was utilized in this study. Our study has demonstrated that RpS13 is of paramount importance for spermatogonial development in the *Drosophila* testis. When RNAi-mediated knockdown of RpS13 was carried out under the control of Bam - Gal4, it resulted in different levels (weak, moderate, and strong) of early-stage germ cell accumulation at the apical tip of the testes (Fig. 1A).

**Fig. 1.**
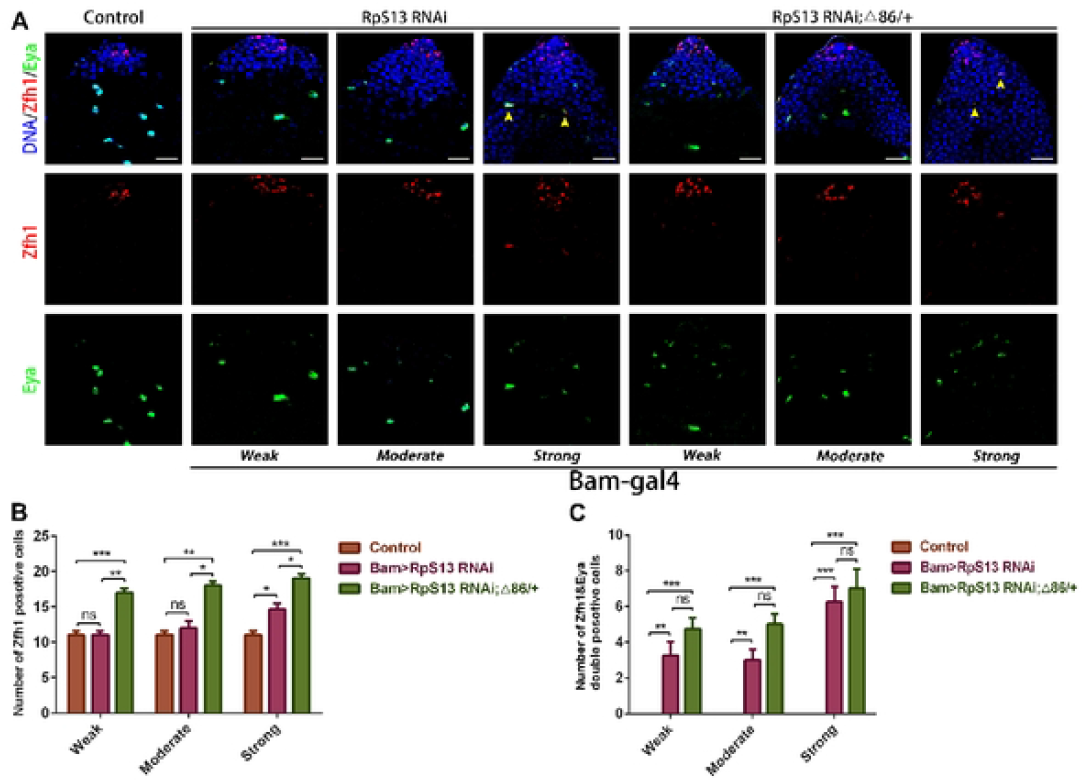
The differentiation of cyst cells was disrupted by reduction of RpS13 in spermatogonia. **A**. Immunostaining of Zfhl (red) and Eya (green) at the apex of W^1118^, Bam>*RpS13* RNAi and Bam>*RpS13* RNAi; Δ86/+ testes. Yellow arrowheads indicate representative Zfhl&Eya-positive cells. **B**. The number of Zfhl positive cells in W^1118^ (n = 3), Bam> *RpS13* RNAi (n = 3) and Bam> *RpS13* RNAi; A86/+ (n = 3) testes. **C**. The number of Zfhl&Eya double positive cells in ^1118^ (n = 3), Bam> *RpS13* RNAi (n = 3) and Bam> *RpS13* RNAi; A86/+ (n = 3) testes. \**P* < 0.05, \*\**P*< 0.01, \*\*\**P*< 0.001, ns represents no significance. Scale bars, 20 μm.

GSC regulators have been demonstrated to play a critical role in maintaining cyst cell survival and preserving testicular architecture [23]. In this study, we employed two well-characterized cyst cell markers, Zfh1 and Eya, to systematically investigate the persistence of cyst cells within the spermatogonial niche following RpS13 depletion. While Eya specifically marked mature cyst cells in control testes, Zfh1 exhibited dual specificity, identifying both CySCs and their early-stage progeny. It is noteworthy that the disruption of Zfh1 serves as an essential regulatory mechanism driving cyst cell lineage progression [27]. Contrary to expectations, quantitative analysis revealed a significant elevation in the population of early-stage and mature cyst cells in both Bam>*RpS13* RNAi and Bam>*RpS13* RNAi; Δ86/+ testes compared to control testes (Fig. 1A-1B and Fig. S1). Particularly striking was the observed increase in Zfh1/Eya double-positive cell populations in these experimental genotypes (Fig. 1C). These cumulative findings suggest that RpS13 depletion in spermatogonia exerts non-cell-autonomous effects that disrupt the developmental trajectory of cyst cells, potentially through altered intercellular signaling pathways within the testicular microenvironment. These results strongly emphasize the indispensable role of RpS13 in regulating the progression and differentiation of spermatogonia.

### Rho1 and RpS13 competitively regulate cyst cell differentiation in *Drosophila* spermatogonia

Our previous study has shown that Rho1 mimics the RpS13 phenotype in the stem cell niche of *Drosophila* testis [23]. Here, we examined the relationship between Rho1 and RpS13 in more detail. Rho1 expression was increased in S2 cells with RpS13 siRNA, as shown in Fig. S2A. On the other hand, the expression of RpS13 was downregulated in Rho1 siRNA S2 cells (Fig. S2B). These results suggest that RpS13 and Rho1 have a mutually inhibitory association throughout *Drosophila* embryo development.

To further investigate the impact of Rho1 on cyst cells, we subsequently performed immunofluorescence staining for Zfh1 and Eya in Bam>*Rho1* RNAi and Bam>*Rho1* RNAi; Δ86/+ *Drosophila* testes. Surprisingly, compared to the control group, the number of Zfh1+ and Eya+ cyst cells was significantly increased in Bam>*Rho1* RNAi and Bam>*Rho1* RNAi; Δ86/+ testes (Fig. 2A-2B and Fig. S3). Simultaneously, the number of double-positive Zfh1/Eya cyst cells was also significantly elevated in Bam>*Rho1* RNAi and Bam>*Rho1* RNAi; Δ86/+ *Drosophila* testes (Fig. 2C). These data suggest that Rho1 may compete with RpS13 to regulate cyst cell differentiation during the TA divisions of spermatogonia.

**Fig. 2.**
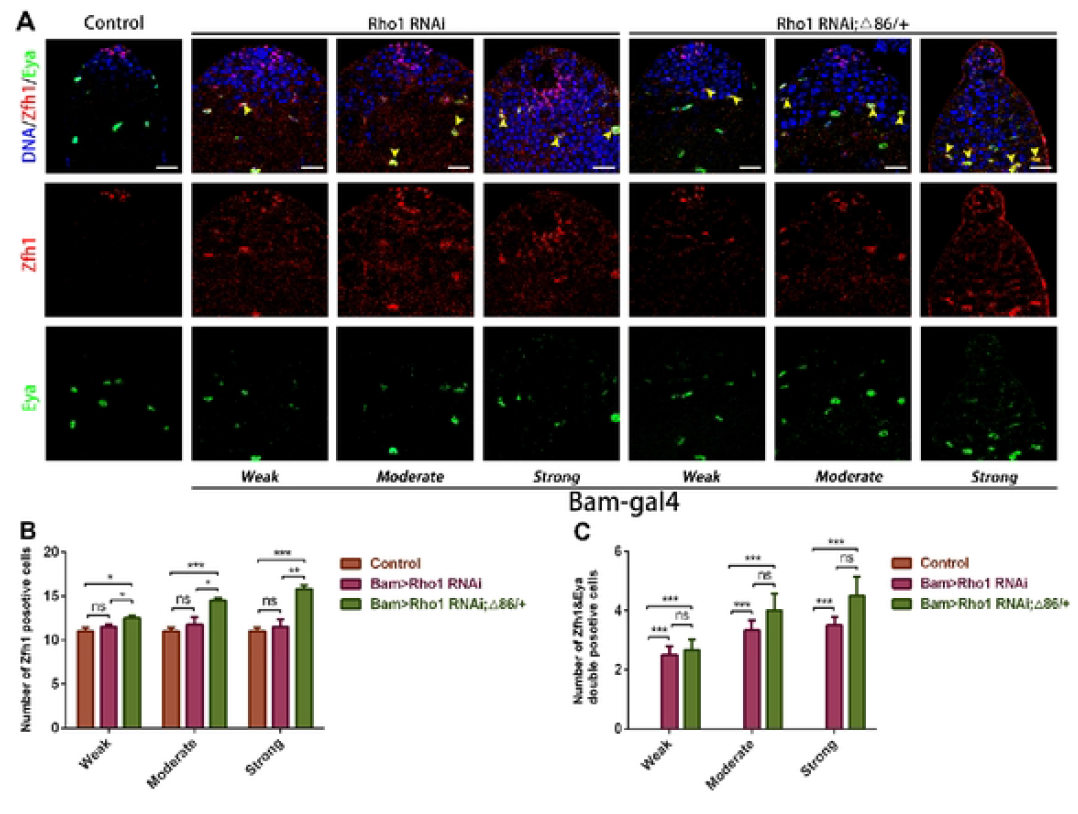
Reduction of Rhol in spermatogonia disrupted cyst cells differentiation. **A**. Immunostaining of Zfhl (red) and Eya (green) at the apex of W1118, Bam>*Rho1* RNAi and Bam>*Rho1* RNAi; Δ86/+ testes. Yellow arrowheads indicate representative Zfhl&Eya-positive cells. **B**. The number of Zfhl positive cells in W^1118^ (n = 3), Bam>*Rho1* RNAi (n = 3) and Bam>*Rho1* RNAi; d86/+ (n = 3) testes. **C**. The number of Zfhl&Eya double positive cells in W^1118^ (n = 3), Bam>*Rho1* RNAi (n = 3) and Bam>*Rho1* RNAi; Δ86/+ (n = 3) testes. \**P* < 0.05, \*\**p* < 0.01, \*\*\**p* < 0.001, ns represents no significance Scale bars, 20 μm.

### Spermatogonial RpS13 controls DE-cad- and Arm-mediated cell adhesion function

Several studies have highlighted the critical role of somatic encapsulation quality in supporting the transplantation and maintenance of spermatogonia. In the *Drosophila* testis, DE-cadherin-mediated cell adhesion junctions are known to play a pivotal role in regulating the stem cell niche [28]. In this study, we investigated the expression pattern of DE-cadherin (DE-cad) under conditions of RpS13 inactivation in spermatogonia. In control testes, DE-cad was predominantly expressed in somatic cyst cells (Fig. 3A). Intriguingly, we observed a progressive increase in DE-cad expression in both Bam>*RpS13* RNAi and Bam>*RpS13* RNAi; Δ86/+ testes, correlating with the severity of cell differentiation impairment (Fig. 3B). In wild-type *Drosophila* testes, Arm is localized along the membranes of both SCCs and hub cells [5]. Similarly, we found that Arm was strongly expressed in somatic cyst cells in control testes. Notably, the expression of Arm was significantly upregulated in Bam>*RpS13* RNAi and Bam>*RpS13* RNAi; Δ86/+ testes, with higher levels corresponding to more severe spermatogonial differentiation defects (Fig. 3C-D). Interestingly, in testes exhibiting pronounced differentiation defects, certain nuclei appeared clustered and displayed reduced levels of DE-cad and Arm compared to surrounding regions (Fig. 3A and D). These findings suggest that RpS13 may modulate cell adhesion junctions during the TA divisions of spermatogonia, potentially influencing their differentiation and niche maintenance.

**Fig. 3.**
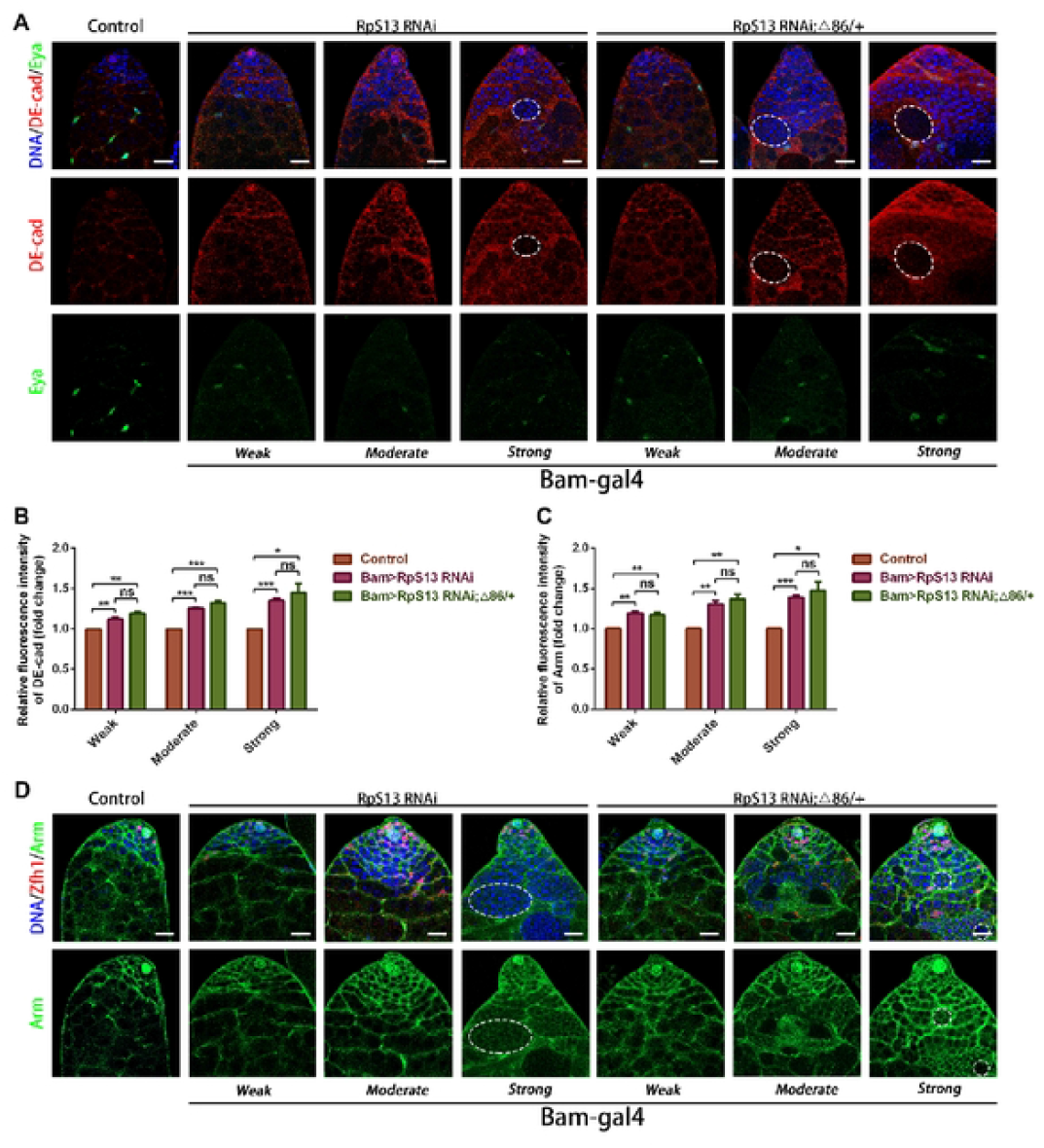
Loss of RpS13 in spermatogonia disrupted cell adhesions. **A**. Immunostaining ofDE-cad (red) and Eya (green) at the apex ofW^1118^, Bam>*RpS13* RNAi and Bam>*RpS13* RNAi; Δ86/+ testes. White circles represent aggregated tumor like cells. B. The relative fluorescence intensity of DE-cad in W ^1118^ (n = 3), Bam>*RpS13* RNAi(n = 3) and Bam>*RpS13* RNAi; Δ86/+ (n = 3) testes. C. The relative fluorescence intensity of Arm in W^1118^ (n = 3), Bam>*RpS13* RNAi (n = 3) and Bam>*RpS13* RNAi; 686/+ (n = 3) testes D. lmmunostaining of ZfuI (red) and Ann (green) at the apex ofW^1118^, Bam>*RpS13* RNAi and Bam>*RpS13* RNAi; Δ86/+ testes. White circles represent aggregated tumor-like cells. \**P* < 0.05, \*\**P* < 0.01, \*\*\**P* < 0.001, ns represents no significance. Scale bars, 20 μm.

### Rho1 regulates adhesion junctions during *Drosophila* spermatogonia TA-divisions

Previous studies have established that the small GTPase Rho1 plays a critical role in regulating cadherin-based adhesion during embryonic morphogenesis and is implicated in the mislocalization of DE-cad protein [29, 30]. In this study, we initially observed that Rho1 was predominantly localized in somatic cyst cells (Fig. S4A). Furthermore, Rho1 protein levels were significantly reduced in both Bam>*Rho1* RNAi and Bam>*Rho1* RNAi; Δ86/+ testes compared to control testes, particularly in nuclear clusters (Fig. S4B). To investigate the impact of Rho1 deficiency on cell adhesion during the TA-division of spermatogonia, we performed immunostaining for DE-cad and Arm. Consistent with our hypothesis, both DE-cad and Arm levels were elevated in Bam>*Rho1* RNAi and Bam>*Rho1* RNAi; Δ86/+ testes relative to controls, with this phenotype being more pronounced in testes exhibiting severe germline differentiation defects (Fig. 4). Notably, nuclear aggregates in Bam>*Rho1* RNAi and Bam>*Rho1* RNAi; Δ86/+ testes displayed markedly lower levels of DE-cad and Arm compared to adjacent regions (Fig. 4A and D). These findings strongly suggest that Rho1 modulates cell adhesion junctions during the TA division of spermatogonia, recapitulating the phenotype observed with RpS13 inactivation, thereby highlighting a conserved regulatory mechanism in this process.

**Fig. 4.**
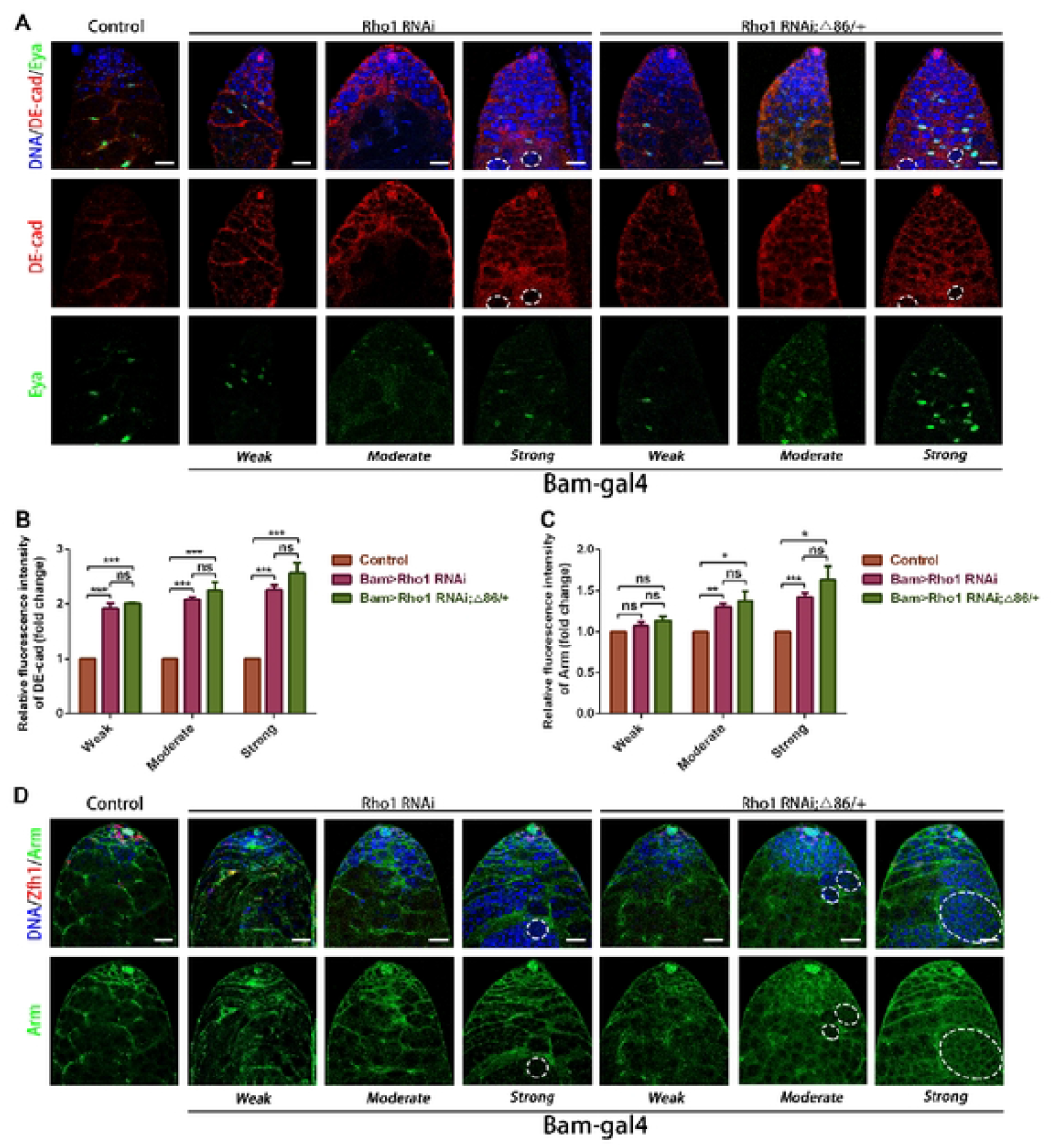
Reduction ofRhol in spermatogonia led to aberrant cell adhesions. **A.** lmmunostaining of DE-cad (red) and Eya (green) at the apex ofW^1118^, Bam>*Rho1* RNAi and Bam>*Rho1* RNAi; Δ86/+ testes. White circles represent aggregated tumor like cells. **B.** The relative fluorescence intensity ofDE-cad in W^1118^, Bam>*Rho1* RNAi and Bam>*Rho1* RNAi; Δ86/+ testes. **C.** The relative fluorescence intensity of Arm in W1118, Bam>*Rho1* RNAi and Bam>*Rho1* RNAi; A86/+ testes. **D.** lmmunostaining of Zfhl (red) and Arm (green) at the apex ofW1118, Bam>*Rho1* RNAi and Bam>*Rho1* RNAi; Δ86/+ testes. White circles represent aggregated tumor-like cells. \**P* < 0.05, \*\**P* < 0.01, \*\*\**P* < 0.001, ns represents no significance. Scale bars, 20 μm.

### RpS13 and Rho1 regulate the expression of dpERK during spermatogonia TA-divisions

ERK (extracellular signal-regulated kinase) signaling is critically involved in regulating the expression and function of putative intercellular adhesion molecules [31]. In *Drosophila* testis, somatic downregulation of ERK activity has been shown to disrupt Arm staining, indicating its essential role in cellular adhesion [5]. To investigate whether RpS13 and Rho1 in spermatogonia modulate cell adhesion junctions through ERK signaling, we examined the expression pattern of dpERK in *Drosophila* testis. In control testes, dpERK was predominantly localized in CySCs and exhibited a slightly patchy distribution (Fig. 5A) [5]. In contrast, dpERK staining displayed a predominantly filamentous pattern in both Bam>*RpS13* RNAi and Bam>*RpS13* RNAi; Δ86/+ testes (Fig. 5A). These findings suggest that RpS13 and Rho1 may influence cell adhesion dynamics by modulating ERK signaling activity in spermatogonia, highlighting a potential mechanistic link between these components and cellular adhesion processes.

**Fig. 5.**
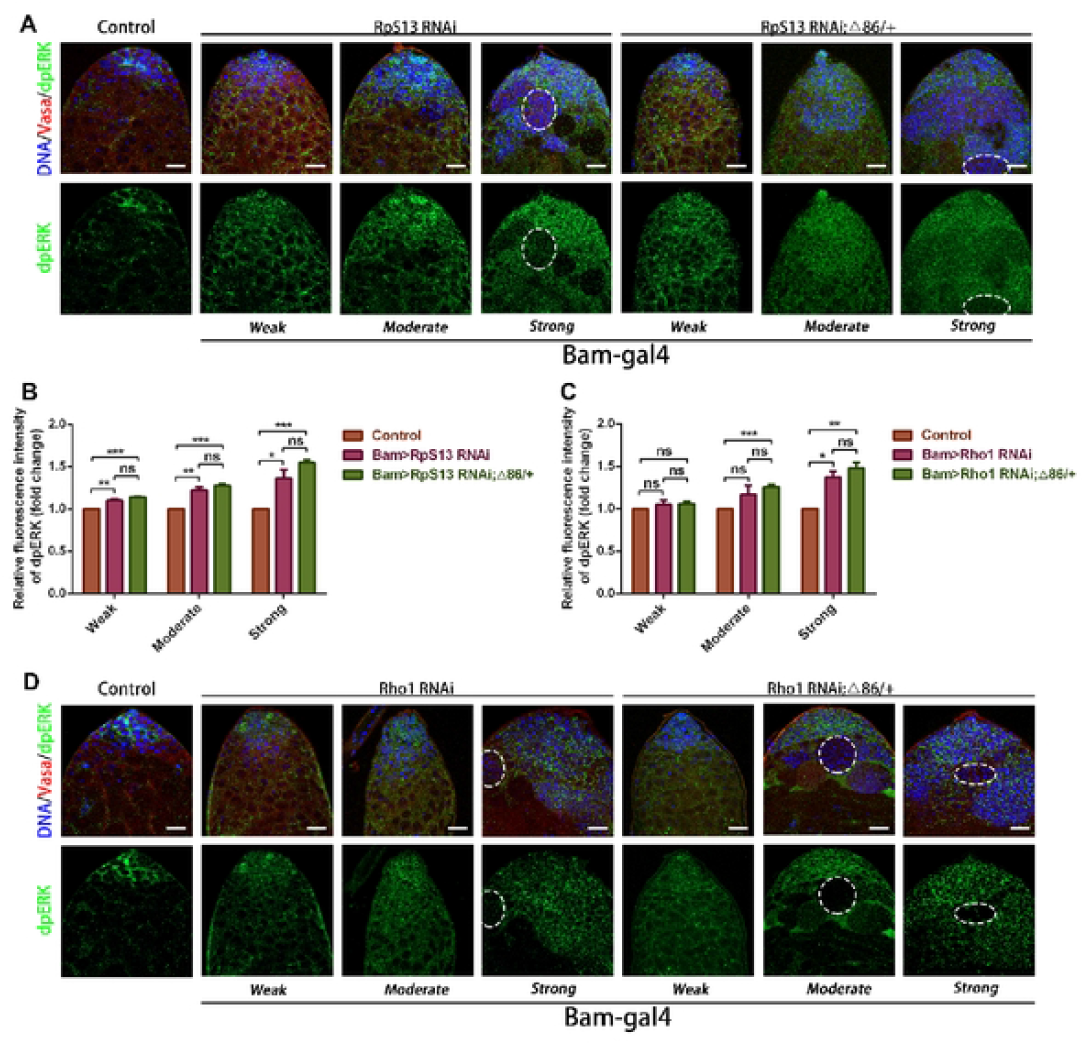
Reduction of RpS13 or Rhol in spermatogonia upregulated dpERK expression. **A**. Immunostaining ofVasa (red) and dpERK (green) at the apex ofW^1118^, Bam>*RpS13* RNAi and Bam>*RpS13* RNAi; Δ86/+ testes. White circles represent aggregated tumor like cells. **B**. The relative fluorescence intensity of dpERKin W^1118^ (n = 3), Bam>*RpS13* RNAi (n = 3) and Bam>*RpS13* RNAi; Δ86/+ (n = 3) testes. **C**. The relative fluorescence intensity of dpERK in W ^1118^, Bam>*Rho1* RNAi and Bam>*Rho1* RNAi; Δ86/+ testes. **D**. Immunostaining ofVasa (red) and dpERK (green) at the apex ofW^1118^, Bam>*Rho1* RNAi and Bam>*Rho1* RNAi; Δ86/+ testes. White circles represent aggregated tumor like cells. \**P* **<** 0.05, \*\**P* < 0.01, \*\*\**P* **<** 0.001, ns represents no significance. Scale bars, 20 μm.

Meanwhile, the expression level of dpERK was found to increase in correlation with the amelioration of spermatogonial differentiation issues (Fig. 5B). Additionally, dpERK levels were significantly lower in the nuclear clusters compared to the surrounding regions. Similar alterations in dpERK staining patterns were observed in the Bam>*Rho1* RNAi and Bam>*Rho1* RNAi; Δ86/+ testes (Fig. 5C-D). These findings suggest that the abnormal activation of dpERK is a consequence of the functional inactivation of RpS13 and Rho1 in spermatogonia. Based on these observations, we propose that *RpS13* RNAi in spermatogonia leads to disruptions in adhesion junctions, which subsequently causes developmental abnormalities in cyst cells. This disruption may, in turn, trigger ERK activation within cyst cells, linking adhesion junction integrity to ERK signaling dynamics in the testis microenvironment.

## Discussion

Research has established that soma-germline signaling plays an instructional role in regulating stem cell fate during spermatogenesis [8, 32]. While cyst cells are not required for the maintenance of GSC-like cells adjacent to the hub, they are essential for germ cells to properly enter the TA program, an early stage of differentiation where stem cell daughters undergo a limited number of mitotic divisions before proceeding to terminal differentiation [33]. These findings underscore the critical importance of SCCs in spermatogonial differentiation. Our research demonstrated that the absence of RpS13 and Rho1 results in the blockage of spermatogonia in the *Drosophila* testis. Additionally, we observed that spermatogonia with reduced RpS13 and Rho1 exhibited an increased number of Zfh1-positive and Zfh1/Eya double-positive cells, indicating impaired cyst cell development. Based on these findings, we conclude that RpS13 and Rho1 non-autonomously regulate cyst cell differentiation, thereby promoting the TA division of spermatogonia in the *Drosophila* testis.

Classical cadherins play a pivotal role in cell signaling and the regulation of cell proliferation and differentiation [34, 35]. The presence of Arm along the cyst contour suggests that adherens junctions may be crucial for the formation and maintenance of somatic encapsulation. One study demonstrated that the loss of Arm in *Drosophila* testis disrupts the somatic permeability barrier surrounding the germinal vesicle cysts, leading to a noticeable reduction in the number of elongated and mature spermatids. This indicates a potential blockade in differentiation beyond the spermatocyte stage [5]. Additionally, several studies have highlighted that Arm can bind to cadherin at adhesion junctions and is localized along the membranes of both somatic cyst cells and hub cells in the *Drosophila* testis, underscoring the essential role of Arm-mediated cell adhesions in the stem cell niche [36-38].

In our present study, we observed that the knockdown of RpS13 and Rho1 in *Drosophila* spermatogonia resulted in an increased expression of DE-cad and Arm. This suggests that RpS13 and Rho1 may regulate the differentiation program of somatic cyst cells through DE-cad- and Arm-mediated adhesion junctions. An alternative hypothesis is that defects in RpS13 and Rho1 lead to an accumulation of undifferentiated somatic cyst cells, thereby promoting heightened expression of DE-cad and Arm at the somatic cell interface. However, these two hypotheses require further experimental validation to establish their validity in the context of somatic cyst cell differentiation and adhesion dynamics.

In this study, we observed that spermatogonia deficient in RpS13 exhibit an aberrant expression pattern of dpERK. ERK activation in SCCs is crucial for synchronizing germ cell divisions within a cyst at each stage of the TA process [5]. A growing body of research highlights the connection between ERK signaling and ribosomal biogenesis during development. Savraj S. Grewal proposed that the regulation of ribosome synthesis serves as a key effector of the Ras/ERK signaling pathway, enabling cells to enhance protein synthesis capacity and thereby promote growth and proliferation [39]. Specifically, ERK has been demonstrated to phosphorylate and activate Mnk, a downstream kinase, which in turn directly phosphorylates eIF4E, a critical translation initiation factor, thereby enhancing its function [40]. In the *Drosophila* testis, eIF4E is essential for spermiogenesis [41, 42]. These findings underscore the interplay between ERK signaling, ribosomal biogenesis, and spermatogenesis in *Drosophila*.

Our experiments unveiled an intriguing phenotype: the absence of RpS13 and Rho1 resulted in elevated levels of DE-cad, Arm, and dpERK in the testis, accompanied by the emergence of nuclear clusters within the mutant testes. Notably, the levels of DE-cad, Arm, and dpERK within these clusters were markedly lower compared to those in the surrounding regions. Consequently, the cells within these clusters exhibited heterogeneity. The ability of cells to segregate into distinct populations is fundamental for the formation of diverse structures during morphogenesis. Steinberg et al. were the first to elucidate the pivotal role of cadherins in cell sorting by proposing the ‘differential adhesion hypothesis’ [43]. In their experiments, cell aggregates expressing varying levels of P-cadherin were co-cultured, revealing that different cell populations eventually separated and organized into a spherical structure. Cells with higher P-cadherin expression accumulated at the core of the sphere, while those with lower expression formed an outer layer, creating a ‘sphere-within-a-sphere’ configuration [44].

Further evidence supporting the role of classical cadherins in mediating cell sorting was demonstrated through studies utilizing *Drosophila* oogenesis as a model system [45, 46]. Importantly, a reduction in surface adhesion molecules in tumor cells may enhance their metastatic potential by facilitating the dissociation of intercellular contacts, thereby promoting diffusion [47, 48]. Based on these findings, we hypothesize that clumps of cells with diminished adhesion protein expression represent a subset of tumor-like cells, and that the absence of RpS13 and Rho1 induces a differential adhesion effect during the TA-divisions of spermatogonia. This differential adhesion could contribute to the observed cellular heterogeneity and cluster formation in the mutant testes.

In summary, the experimental results presented in this study demonstrate that RpS13 is intricately linked to Rho1-mediated adhesion signaling in the *Drosophila* testis and plays a non-autonomous role in regulating the differentiation of cyst cells during spermatogonia TA-divisions. Our findings provide robust evidence that RpS13 modulates germline differentiation through soma-germline adhesion junctions. This research establishes a foundational framework for exploring the intricate relationships between germline and somatic cells, while also broadening our comprehension of the molecular mechanisms governing the regulation of spermatogonia TA-divisions. These insights contribute significantly to the field of developmental biology and highlight the importance of adhesion signaling in germline-soma communication during spermatogenesis.

## Acknowledgement

The study was supported by National Natural Science Foundation of China (82072088).

## Declaration of competing interest

The authors declare that they have no conflicts of interest.

## Ethical statement

This study was approved by the Ethics Committee of Yangzhou University.

## Author contributions

Min Wang and Lingling Gao initiated the project and designed the research; Min Wang, Li Zhang and Jianbo Xu performed most of the experiments, Yanxin Zhang, Min Jiang, Jingzhe Song and Xilong Liu performed data collection and analysis. Min Wang wrote the manuscript with the assistance of Dan Lu. All authors read and approved the final manuscript.

## Data availability statement

The data that support the findings of this study are available from the corresponding author upon reasonable request.

## Notes

### Competing Interest Statement

The authors have declared no competing interest.

